# Reorganization of corticospinal projections after prominent recovery of finger dexterity from partial spinal cord injury in macaque monkeys

**DOI:** 10.1101/2023.02.19.529046

**Authors:** Masahiro Sawada, Kimika Yoshino-Saito, Taihei Ninomiya, Takao Oishi, Toshihide Yamashita, Hirotaka Onoe, Masahiko Takada, Yukio Nishimura, Tadashi Isa

## Abstract

We investigated morphological changes in the corticospinal tract (CST) to understand the mechanism underlying recovery of hand function after lesion of the CST at the C4/C5 border in 7 macaque monkeys. All monkeys exhibited complete recovery of precision grip success ratio within a few months. The trajectories and terminals of CST from the contralesional (n = 4) and ipsilesional (n = 3) hand area of primary motor cortex (M1) were investigated at 5-29 months after the injury using an anterograde neural tracer, biotinylated dextran amine (BDA). Reorganization of the CST was assessed by counting the number of BDA-labeled axons and button-like swellings in the gray and white matters. Rostral to the lesion (at C3), the number of axon collaterals of the descending axons from both contralesional and ipsilesional M1 entering the ipsilesional and contralesional gray matter, respectively, were increased. Caudal to the lesion (at C8), axons originating from the contralesional M1, descending in the preserved gray matter around the lesion, and terminating in ipsilesional laminae VI/VII and IX were observed. In addition, axons and terminals from the ipsilesional M1 increased in the ipsilesional laminae VI/VII and IX after recrossing the midline, which were not observed in intact monkeys. Conversely, axons originating from the ipsilesional M1 and directed toward the contralesional laminae VI/VII and IX decreased. These results suggest that multiple reorganizations of the corticospinal projections to spinal segments both rostral and caudal to the lesion originating from bilateral M1 underlie a prominent recovery in long-term after spinal cord injury.

**SIGNIFICANCE STATEMENT:** Previous studies have shown that dexterous finger movements recover prominently after lesion of the corticospinal tract (CST) at the mid-cervical segments through rehabilitative training in macaque monkeys. Here, we show reorganization of the CST including sprouting of axons originating from the contralesional and ipsilesional motor cortex in the gray matter both caudal and rostral to the lesion, including a re-direction of the CST to hand motoneurons in the monkeys 5-29 months after the lesion. Thus, multiple mechanisms of reorganization of CST axons underlie the recovery of impaired cortico-motoneuronal pathways for the long-term recovery of finger dexterity.

## INTRODUCTION

Hand use is a critical component in our daily life, and recovery of hand function is a major issue for patients with neuronal damage such as spinal cord injury (SCI) (Anderson 2004). Investigating the neuroanatomical basis of recovery in an animal model that shows prominent recovery of finger dexterity is critical for future development of therapeutic strategies to promote the recovery of hand functions. However, the recovery of finger dexterity has been shown to be difficult after large injuries such as hemisection (Galea and Darian-Smith 1997; Rosenzweig et al. 2010) or subhemisection without drug treatment (Nakagawa et al. 2015). In this regard, it should be noted that macaque monkeys with lesions limited to the dorsolateral funiculus (DLF) which transected the lateral corticospinal tract (l-CST) and rubrospinal tract show prominent recovery of precision grip in several weeks after injury (Sasaki et al. 2004), and our laboratory has been investigating the mechanisms of recovery in this model (Isa, 2017, 2019).

Direct cortioco-motoneuronal connections, i.e., the monosynaptic pathway from corticospinal neurons to spinal motoneurons, play a pivotal role in the control of dexterous finger movements in primates (Lawrence and Kuypers, 1968; Kuypers, 1982; Lemon, 2008). Conversely, our previous series of reports documented that indirect cortico-motoneuronal pathways mediated by propriospinal neurons (PNs) located in spinal segments rostral to motoneurons exist in primates, are connected to motoneurons, and partially contribute to the control of dexterous finger movements (Alstermark et al., 1999; Isa et al., 2006; Alstermark et al. 2011; Kinoshita et al., 2012). Studies on the C4/C5 DLF lesion model demonstrated that the indirect cortico-motoneuronal connections via C3-C4 PNs mediating the command for dexterous finger movements were strengthened and contributed to the prominent recovery of dexterous finger control (Sasaki et al. 2004; Tohyama et al., 2017). Functional reorganization has also been demonstrated at the cortical level. After monkeys showed full recovery of finger dexterity from the C4/C5 DLF lesion, multiple motor-related cortical areas on bilateral sides which are origins of CST showed increased activity (Nishimura et al., 2007; Sawada et al., 2015; Suzuki et al., 2020). These observations suggest that plastic changes in the corticospinal connections from bilateral motor cortices are involved in prominent functional recovery, however it remains unclear whether these functional changes are accompanied by morphological changes in the corticospinal projections.

Previous studies revealed that the CST exhibits substantial sprouting both caudal and rostral to the lesion after the recovery of hand function in the hemisection/subhemisection models of SCI (Fouad et al., 2001; Freund et al. 2006; Rosenzweig et al., 2010; Nakagawa et al., 2015). However, in these models, the recovery of dexterity was limited due to the large lesion of the spinal cord. Therefore, a quantitative analysis of the reorganization of CST in the partial SCI model with prominent recovery and comparison with the results of hemisection/subhemisection model with poor recovery may be valuable for understanding the basis of prominent recovery of dexterous finger movements. For this purpose, we investigated quantitatively the distribution of CST axons and their terminals in the white matter and each lamina of the gray matter rostral and caudal to the lesion limited to the DLF at the C4/C5 border in macaque monkeys that showed prominent recovery of dexterous finger movements.

## MATERIALS AND METHODS

### Statement of Ethics

All applicable institutional and governmental regulations concerning the ethical use of animals were followed during the course of this research. The experiments were subjected to prior reviews by the ethical committee of the National Institutes of Natural Science and were performed in accordance with the guideline of the Ministry of Education, Culture, Sports, Science, and Technology (MEXT) of Japan and NIH guidelines of the Care and Use of Laboratory animals.

### Animals

Seven male macaque monkeys (macaca fuscata; Di, To, Al, Gr, De, As, Fu, ranging from 3.6 kg to 8.6 kg) (Table 1) were used in this study. To investigate the anatomical changes induced during functional recovery after SCI, the data obtained from the monkeys with SCI were compared with data from the five non-lesioned control monkeys (Mo-1 (male), 2 (female), 3 (female), 4 (female), 5 (male)) reported in our previous study (Yoshino-Saito et al., 2010).

**Table 1.** Experimental time course and injection sites of BDA in each monkey whose data were used in this manuscript.

| Name | Injected side | Side to lesion | Days to injection | Survival time after injection | Total survival time | N of tracks | N of injection sites | Injection depths (mm) (or) | Injected amount (μL/site) |
| --- | --- | --- | --- | --- | --- | --- | --- | --- | --- |
| Di | Right | Contra | 155 | 75 | 230 | 6 | 15 | 1.5, 3.0<br>4.5 | 0.5 |
| To | Right | Contra | 563 | 114 | 677 | 8 | 20 | 1.5, 3.0<br>4.5 | 0.5 |
| Al | Right | Contra | 278 | 93 | 371 | 7 | 18 | 1.5, 3.0<br>4.5 | 0.5 |
| Gr | Right | Contra | 277 | 178 | 455 | 6 | 12 | 1.5, 3.0<br>4.5 | 0.5 |
| De | Left | Ipsi | 107 | 42 | 149 | 4 | 8 | 3.5, 5.0 | 0.5 |
| As | Left | Ipsi | 124 | 30 | 154 | 4 | 8 | 3.5, 5.0 | 0.5 |
| Fu | Left | Ipsi | 731 | 139 | 870 | 7 | 18 | 2.0, 5.0<br>8.0 | 0.5 |
| Mo1 | Right |  |  | 104 |  | 5 | 15 | 2.0, 4.0<br>6.0 |  |
| Mo2 | Right |  |  | 103 |  | 5 | 15 | 2.0, 4.0<br>6.0 |  |
| Mo3 | Right |  |  | 80 |  | 4 | 12 | 2.0, 4.0<br>6.0 |  |
| Mo4 | Right |  |  | 92 |  | 7 | 21 | 2.0, 4.0<br>6.0 |  |
| Mo5 | Right |  |  | 94 |  | 5 | 15 | 2.0, 4.0<br>6.0 |  |

### Behavioral training and functional assay of finger dexterity

Monkeys were trained to perform reach and grasp task with their left arms (Fig. 1A). They were trained to grasp pieces of sweet potato (5 mm × 5 mm × 5 mm) through a narrow slit (10 mm in width) on an acrylic panel placed in front of the home cage (Monkeys Di, Al, and Gr) or in front of the monkey chair on which they were seated (Monkeys To, De, As, and Fu) (Fig. 1A). To evaluate the recovery course of finger dexterity, hand movements were filmed during task performance from the radial side (30 frames per second; shutter speed, 1/250 sec). A ‘successful precision grip’ was defined as the monkey grasping the piece of food using only the ventral sides of the index finger and thumb (Fig. 1A). The success rate of precision grip in the 30 trials from beginning of each daily session was calculated throughout the experimental period (Fig. 1B).

**Figure 1.**
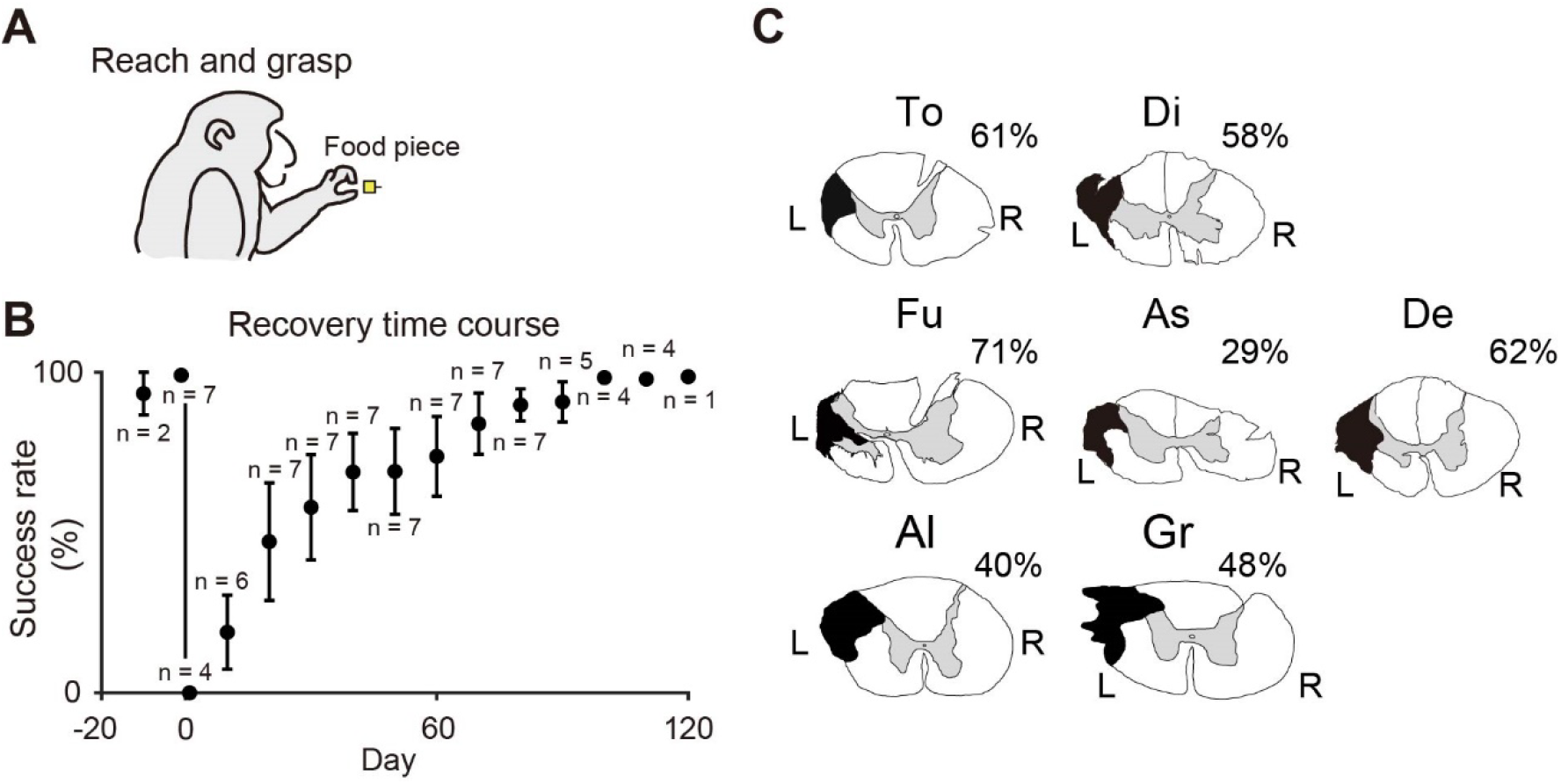
Recovery course of precision grip and spinal cord lesion. **A**. Reach and grasping task (left). Representative image of Precision grip (right). **B**. Recovery time course of the success rate of precision grip in 7 monkeys. Each plot represents the mean success rate. Error bars indicate S.E.M. ‘n’ means the number of monkeys on each day. A successful trial was defined as that in which the monkey succeeded in grasping the food morsel with precision grip using just the pads of the index finger and thumb and bringing it to the mouth to eat without dropping it. Data from the last behavioral test before the SCI (Day -1 or Day 0) are plotted on Day 0. **C**. The extent of spinal cord injury in each animal. The black shaded area indicates the maximum lesion extent of the spinal cord at the border of C4 and C5. The lesion extents of Monkeys Al and Gr were reconstructed from parasagittal sections of their spinal cord. The percentage value beside each panel shows the relative extent of spinal cord injury in each animal (see Methods). L: left, R: right.

### Surgeries

All surgical procedures were performed under sterilized conditions. The monkeys were sedated by intramuscular injection of ketamine (Daiichi-Sankyo, Japan; 10 mg/kg i.m.) plus xylazine (Bayer, Japan; 1 mg/kg i.m.). Atropine (Tanabe Pharma, Japan; 0.01 mg/kg i.m.), ampicillin (Meiji Pharma, Japan; 40 mg/kg i.m.) and dexamethasone (Banyu Pharma, Japan; 0.25 mg/kg) were injected intramuscularly as premedication. During the surgery, anesthesia was maintained with intravenous injection of sodium pentobarbital (Abbott Laboratories, US; 25 mg/kg i.v.) or isoflurane (MSD Animal Health, Japan; 1–2%) inhalation. ECG, SpO2, end-expiratory carbon dioxide pressure, and body temperature were monitored carefully during the surgery. Drips of Ringer’s solution were given throughout surgery. Dexamethasone (NICHI-IKO, Japan; 0.01 mg/kg i.m.), ampicillin (Fujifilm Wako Chemical Corp., Japan; 40 mg/kg i.m.), and ketoprofen (Kissei Pharmaceutical Co., Ltd., Japan; 2.0 mg/kg i.m.) were administered after surgery.

#### Surgery 1: Lesion of the corticospinal tract (CST)

The DLF, where the majority of CST and rubrospinal tract axons course, was transected at the border between the C4 and C5 segments on the left side of the spinal cord, as described previously (Sasaki et al. 2004; Nishimura et al., 2007; Sugiyama et al., 2013) (Fig. 1C). A horizontal strip oriented in a mediolateral direction relative to the DLF was made by inserting a blunt L-shaped hook with a maximum insertion of 5 mm, which corresponds to the distance from lateral convexity of the spinal cord to the lateral edge of gray matter. Then, by using fine forceps, the dorsal part of the DLF was transected from the dorsal root entry zone ventrally to the level of the horizontal strip described above. Finally, the lesion was extended ventrally to the most lateral part of the lateral funiculus using forceps. The opening of the dura mater was closed, and the skin and back muscles were sutured with nylon or silk threads.

#### Surgery 2: Neural tracer injection into the motor cortex

After recording the behavioral data after the CST lesion (Di, 155 days; To, 563 days; Al, 278 days; Gr, 277 days; De, 107 days; As, 124 days; Fu, 731 days) anterograde neural tracer was injected into the M1. Under deep anesthesia with isoflurane, the animal was mounted on a stereotaxic frame. Craniotomy was performed to expose the hand area of M1 on the right (contralesional) side (Di, To, Al, and Gr) or left (ipsilesional) side (De, As and Fu) (Table 1; Fig. 2A and C). Cortical surface was exposed after opening the dura. As we described previously (Yoshino-Saito et al., 2010), 0.5 μl solution of biotinylated dextran amine [BDA; Molecular Probes, Eugene, OR, USA; 10,000 MW; 10% dissolved in 0.01% phosphate buffer (pH7.3)] was injected at each injection site in the hand area of M1 using a 10 μl Hamilton microsyringe. One to three injections were made in each track and totally 6-8 tracks were injected with BDA in each animal (details are shown in Table 1).

**Figure 2.**
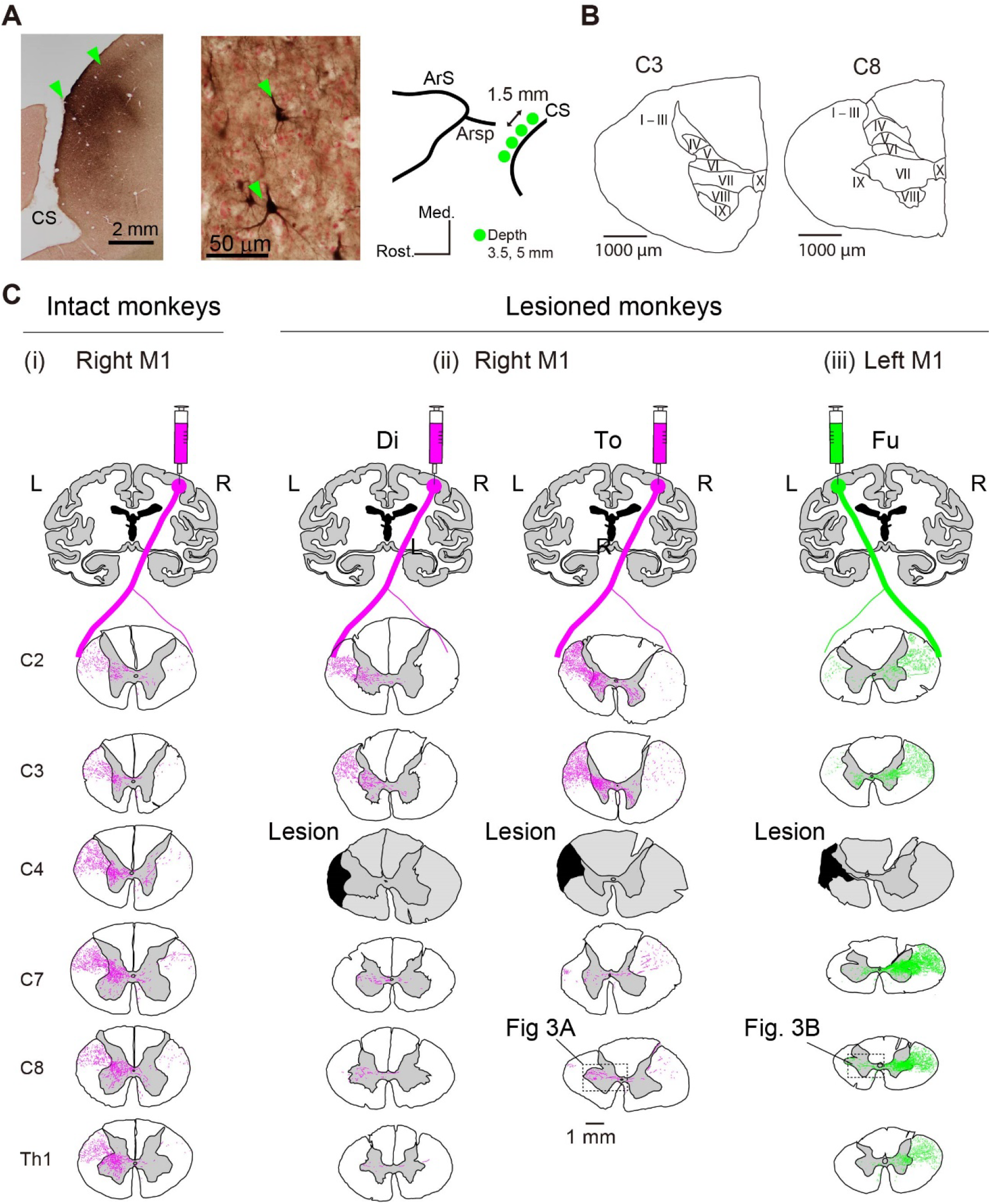
Injection sites of BDA and distribution of the BDA-labeled axons at different segments from C2 to Th1. **A. Left:** Example of injection sites in M1; bulk of BDA-reaction product at the injection site in the gray matter of the rostral bank of the central sulcus (CS). **Center:** A high-magnification view of BDA-labeled pyramidal neurons in layer V at the injection site in Mo Fu. Scale bars in left and center panel indicate 2 mm and 50 μm respectively. **Right:** Schematic distribution of injection tracks of BDA (Green area). Detailed location of injection sites is shown in Table 1. Arc, arcuate sulcus; Arsp, Arcuate sulcus spur. **B**. Definition of the laminae of Rexed in the C3 and C8 spinal segments (Rexed, 1952). Scale bars indicate 1 mm. **C**. Distribution of BDA-labeled axons in the C2, C3, C4, C7, C8, and Th1 spinal segments originating from the right M1 in the intact monkey (Mo-4 from Yoshino-Saito et al. (2010)) (D(i)), and from the contralesional (right) M1 in Monkey To (D(ii)) and from the ipsilesional (left) M1 in Monkey Fu (D(iii)). The black shaded area in the panels of C4 in D(ii) and D(iii) indicates the lesion extent at the border of C4 and C5 in these animals Scale bar = 1 mm.

According to our previous study (Nishimura et al. 2007), the locations of the digit region of M1, as indicated by twitches of the digits and wrist induced by train stimuli of 15 pulses (biphasic pulses of 0.1 ms cathodal and 0.1 ms anodal, at 333 Hz) with the threshold below 40 μA (often below 10 μA) under sedation with ketamine (Daiichi-Sankyo, Japan, 10 mg/kg, i.m.), were distributed between 11 and 17 mm lateral from the midline and 1–2 mm anterior to the central sulcus (CS) (green shaded area in the right panel of Fig. 2A). The locations of the injection tracks were determined by this information. Injection tracks were in two rows, one was 1 mm anterior to the CS and the other was 2-3 mm anterior to the CS. The former tracks were located along the bank of the CS, which corresponds to the “new M1”, and the latter tracks were located on the convexity of the cortical surface, which correspond to the “old M1”, according to Rathelot and Strick (2009). The injection tracks in each row were separated by 2-3 mm medio-laterally.

### Histological processing

One to six months after the injection of BDA (Di, 75 days; To, 114 days; Al, 93 days; Gr, 178 days; De, 42 days; As, 30 days; Fu, 139 days), monkeys were anesthetized deeply with an overdose of sodium pentobarbital (50-100 mg/kg i.v.) and perfused transcardially with 0.1 M phosphate-buffered saline (PBS, pH 7.3) containing heparin (1 unit/ml), followed by 4% paraformaldehyde in 0.1 M PBS (pH 7.3). Therefore, the total survival periods of the monkeys after the spinal cord injury were 230 days for Di, 677 days for To, 371 days for Al, 455 days for Gr, 149 days for De, 154 days for As, and 870 days for Fu (Table 1). After perfusion, the brain and spinal cord were removed and postfixed in the same fresh PBS containing 2% paraformaldehyde, followed by 10%, 20%, and 30% sucrose. After saturation with sucrose, a series of 50 μm thick sections of the brain, including the injection sites, and cervical and thoracic spinal cord from C2 to Th2 segments were obtained using a freezing microtome (Thermo Scientific, USA). The spinal segments were determined by identifying the level of dorsal and ventral rootlets. In 5 monkeys (Di, To, Fu, De, and As), the spinal cord was sectioned coronally, while in 2 monkeys (Al and Gr), the spinal cord was sectioned longitudinally. After BDA-labeling, the sections were mounted on gelatin-coated glass slides. The sections were collected serially in 0.1 M PBS, pH 7.4. Some sections were Nissl-stained with 1% Cresyl Violet to estimate the relative extent of the spinal cord injury. Following the methods described previously (Sugiyama et al. 2013), the extent of lesion (percent) in the DLF was determined by subtracting the ratio of remaining area of the lateral and ventral funiculi on the lesioned side divided by the whole area of the lateral and ventral funiculi from 100%.

In order to visualize the BDA labeling, the remaining sections were processed using the avidin-biotin-peroxidase method (Vector, US) with 3,3’-diaminobenzidine (DAB; Sigma, USA) as the chromogen. Sections were washed twice in 0.05 M PBS for 10 min, followed by incubation with 0.6% hydrogen peroxide in methanol for 30 min to reduce the activity of endogenous peroxidase. After reduction, sections were washed 3 times in PBS, and incubated for 120 min in a solution containing 50 fold-diluted ABC in 0.05 M PBS plus 0.4% Triton X-100. The sections were rinsed twice in PBS and twice in 0.05 M Tris-buffered saline. The peroxidase was reacted and visualized by incubating the sections in a solution containing 0.01% DAB, 1.0% nickel ammonium sulfate, and 0.0003% hydrogen peroxide in 0.05 M Tris buffer, for 10–15 min at room temperature. The reaction was terminated by rinsing in Tris buffer followed by two rinses in PBS. After the DAB-nickel staining, the sections were mounted on gelatin-coated glass slides, some of which were lightly counterstained with 1% Cresyl Violet, dehydrated, and coverslipped.

### Analysis in coronal sections

We chose the C3 and C8 segments for quantitative analysis of the CST axons and terminals as the examples of segments rostral and caudal to the lesion, respectively, because the C3 segment contains a large fraction of the cell bodies of PNs, which presumably mediate the CST inputs to motoneurons, and the C8 segment contains the spinal motoneurons innervating the hand muscle and their premotor interneurons. Therefore, we expected to observe plastic changes in the CST at these segments. In order to quantify the number of axons and terminal button-like swellings, three sections separated by 400 μm intervals were selected from each segment. Detailed trajectories of BDA-labeled axons and their terminals were traced along with a section outline at 10x magnification, using a camera lucida attached to a light microscope (Nikon, Japan).

We traced and counted the number of BDA-labeled axons separately in the white matter including the dorsolateral, ventromedial, and dorsal funiculi on both sides to the BDA injections in the M1. All the axons, including those that ran in the longitudinal and transverse direction relative to the rostrocaudal axis of the spinal cord, were counted. We defined axons whose length was less than 150 μm as longitudinal axons, and those whose length was more than 150 μm as traverse axons in the coronal section, respectively, similar to our previous study (Yoshino-Saito et al. 2010). To evaluate the number of collateral branches rostral to the lesion, we divided the number of transverse axons (>150 μm) by the total number of axons (including longitudinal and transverse axons) in the white matter of bilateral DLF of the C3 segment. We compared these ‘transverse ratios’ of the lesioned monkeys with those of the intact monkeys.

The number of axons labeled with BDA was counted in each lamina of the gray matter, regardless of the length included in each section, on both the ipsilateral and contralateral sides to the BDA injections. The border of each lamina was defined based on the cytoarchitecture according to the definition by Rexed (Rexed, 1952)(Fig. 2B). When the same axon crossed the border between two laminae, the axon was counted as being included in both laminae. The number of BDA-labeled button-like swellings (see Fig. 5H) were also counted in each lamina in the C3 and C8 segments on both the ipsilateral and contralateral sides to the BDA injections. Then, we divided the number of labeled axons and button-like swellings in each lamina by the total number of the axons in the bilateral C3 DLF at each spinal segment for normalization, considering individual differences of the total number of labeled axons. We compared these values of lesioned monkeys with those of intact monkeys to evaluate the difference in the axonal and terminal distribution between these two groups.

### Analysis in longitudinal sections

In the longitudinal sections, we also evaluated the frequency of collateral branches in the C3 segment as representing the segments rostral to the lesion. At the C3 segment, we counted the number of labeled axons that crossed the border between the white matter and gray matter transversely. Then, we calculated ‘transverse ratios’ similarly to the coronal sections.

To evaluate the frequency of the collateral projections that descended in the gray matter caudal to the lesion, we counted the number of labeled axons within 5 mm caudally from the level of the spinal lesioning. We also measured the direction of axons against the orientation of the central canal, and the length of axons in three longitudinal sections. They were then compared between intact and lesioned monkeys (Fig. 5C-F). Labeled axons were traced by Illustrator 25.2.3 (Adobe, US) and saved in a Scalable Vector Graphics file (Fig. 5C). Then, starting and ending points of the traced axons were determined using the original Matlab script (The MathWorks, Inc., US). The length of axons was defined as the length between the starting and ending points, and the angle of axons was defined as the angle of the line that connects these two points of the axon against the orientation of the central canal. The angle ranges from 0° to 90° (Fig. 5D). The relationship of the length and angle was plotted in Fig. 5E. Based on the scatter plot, a two-dimensional histogram was created for the length and angle of axons, where each bin was normalized by the bin of the axons with the shortest length and highest angle (Fig. 5F). We made the same measurements in the corresponding segments of intact monkeys and compared the results with those of the monkeys with SCI.

### Experimental design and statistical analyses

Statistical analysis was performed using MATLAB. Two sample t-test was performed to compare the percentage of axons and button-like swelling between lesioned and intact monkeys. P values are provided in each figure.

## Results

### Lesion extent

Figure 1C shows the extent of the C4/C5 DLF lesion in the 7 monkeys. The lesions in the monkeys were intended to encompass the normal distribution of the l-CST. The lesion extent ranged from 29% to 71% of the lateral and ventral funiculi, which mostly corresponds with the lesion extent reported previously (Nishimura et al. 2007, 2009; Sugiyama et al. 2013; Sawada et al. 2015; Chao et al. 2019). Lesion extent was regarded as covering at least the majority of the l-CST axons in all monkeys.

### Recovery time course

The success ratio of the precision grip task was nearly 100% before the lesion, but dropped to 0% immediately after the SCI. As shown in Figure 1B, however, the success ratio gradually recovered and reached 100% within 3 months in all 7 monkeys (Fig. 1B), and remained constant afterwards, similar to the results in our previous studies (Nishimura et al. 2007, 2009; Sugiyama et al. 2013; Sawada et al. 2015; Chao et al. 2019).

### Overview of the CST originating from the contralesional (right) and ipsilesional (left) M1

We investigated the trajectories of the CST axons from the contralesional (n = 4) and ipsilesional (n = 3) hand area of M1 using an anterograde neural tracer, BDA (Fig. 2). Figure 2A (left) shows a section along the rostral bank of the central sulcus. As described in the Materials and Methods, multiple injections were made along these tracks (Fig. 2A, right). As shown in this figure, the injection of BDA was successful in labeling cortical neurons located at various depths in the precentral gyrus. The labeled neurons around the injection tracks included large pyramidal neurons, which may include CST neurons in lamina V of M1 (Fig. 2A, middle).

Figure 2C (i) shows the trajectories of CST axons in an intact animal (Mo-4), taken from our previous study (Yoshino-Saito et al. 2010). As described previously, the majority of CST axons (85-98%) descend in the DLF on the contralateral side (left; L) to the injection (right, R) and issue collaterals to various segments in the cervical spinal cord (Kuypers 1982; He et al. 1993; Armand et al. 1997; Rosenzweig et al. 2009). In the upper ∼ middle cervical segments (C2, C3, C4), these collaterals terminate mainly in the contralateral laminae VI and VII (cf. Fig. 2B) and partly to the ipsilateral lamina VIII after recrossing the midline. A small number of CST axons descended in the ipsilateral DLF. Some collaterals appeared to enter the gray matter (see the C4 segment) and terminate in lamina VI, VII and VIII of the ipsilateral side, but it was difficult to follow the axonal trajectory completely because they were mixed with the recrossing axons from the axons descending in the contralateral side. In the lower cervical segments (C7, C8, Th1), in addition to the terminals in laminae VI and VII, the axons reached lamina IX on the contralateral side. Projection to the ipsilateral lamina VIII after recrossing could be observed. A small number of collaterals from the ipsilateral, uncrossed CST in the ipsilateral DLF entered the gray matter. However, the number appeared to be small and their trajectories were difficult to follow (Fig. 2C(i)).

#### CST originating from the contralesional (right) M1

Figure 2C (ii) shows examples of the axonal trajectories of the CST originating from the contralesional (right) M1 in the lesioned animals (Monkeys Di (230 days after SCI) and To (677 days after SCI)). Few labeled axons were observed immediately caudal to the lesion in the ipsilesional DLF in these contralesional-injection group, which suggests that some CST axons remained uncut, but the regrowth of axons penetrating the scar of lesion was minimal, if any. In both monkeys, rostral to the C4/C5 lesion (C2, C3), a number of collaterals issued from the main axons into the left DLF to the gray matter. The majority terminated in lamina VI/VII in the left spinal gray matter, but some crossed the midline and terminated mainly in lamina VIII on the contralesional side. The majority of descending axons in the DLF were transected by the lesion at C4/C5. However, caudal to the lesion (C7, C8), some axons could be observed in laminae VII and IX on the ipsilesional side (“C8” in Fig. 2C (ii)). As shown in the photomicrograph in Figure 3A, these axons are mainly penetrating the section perpendicularly because their horizontal length in the section was shorter than 150 μm. The origin of these axons will be discussed in the next section.

**Figure 3.**
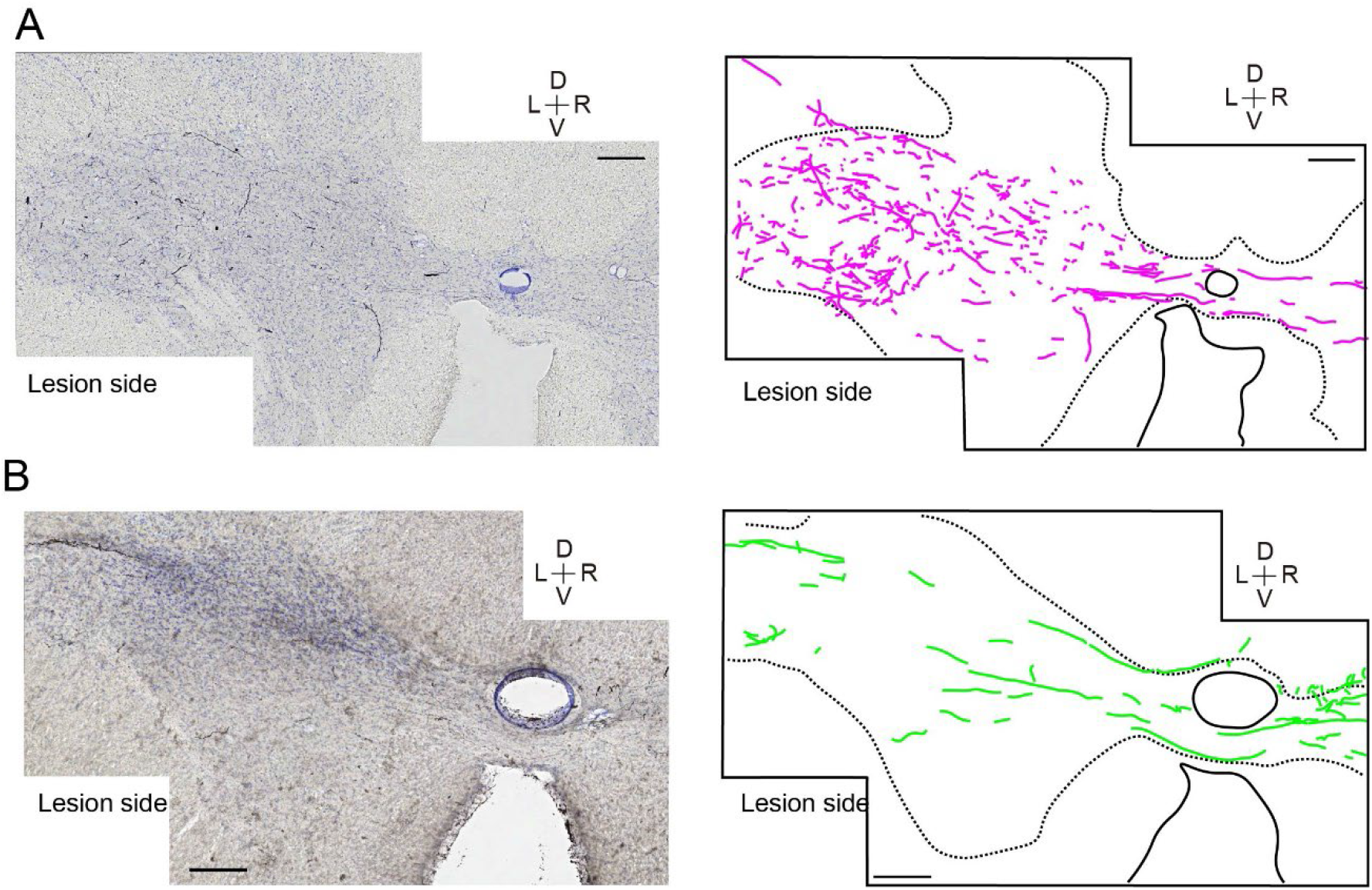
A. CST axons descending from the contralesional M1 in Monkey To at the C8 segment (cf. Fig. 2C(ii)). Photomicrograph in the left and camera lucida tracing of the BDA-labeled axons (in magenta) in the right panel. D, dorsal; V, ventral; L, left; R, right. Scale bar = 200 μm. **B**. CST axons descending from the ipsilesional M1 in Monkey Fu at C8 segment (cf. Fig. 2C(iii)). Photomicrograph in the left and camera lucida tracing of the BDA-labeled axons (in green) in the right panel. Scale bar = 200 μm. Note that the CST axons in **A** are mostly short and penetrate the section perpendicularly, while those in **B** are traversing the section horizontally for a long distance.

#### CST originating from the ipsilesional (left) M1

Figure 2C (iii) shows an example of the axonal trajectories of the CST originating from the ipsilesional (left) M1 in the lesioned animal (Monkey Fu). Rostral to the C4/C5 lesion (C2, C3 segments), the CST axons were mainly located in the contralesional DLF. A number of collaterals issued to laminar VI and VII in the gray matter on the contralesional side at the C3, some of which further crossed the midline and terminated in lamina VIII of the ipsilesional side. Caudal to the lesion (C7, C8, Th1), a large number of collaterals issued from the main axons into the DLF to the gray matter and terminated in laminae VI and VII on the right side. As shown in the photomicrograph in Figure 3B, a majority of these collaterals crossed the midline, traversed the gray matter, and terminated in the ventral horn including lamina IX on the ipsilesional (left) side of the spinal cord.

Table 2 shows the number of BDA-labeled axons in the DLF at the C3 and C8 segments on the ipsilesional and contralesional sides. As shown in the table, the BDA-labeled axons were markedly reduced at the C8 segment in the ipsilesional DLF in the lesioned monkeys after BDA injection in both the contralesional (Di and To) and ipsilesional M1 (Fu, De, As). These results suggest that the transection of CST axons was mostly perfect. Conversely, there was only slight decrease in the number of axons (Mo-2, 3, 4) or even increase (Mo-1) in the number of BDA-labeled axons at the C8 segment compared with the C3 in the intact monkeys. The increase might reflect the branching of the CST axons in the white matter before innervating the gray matter of the lower cervical segments.

**Table 2.**
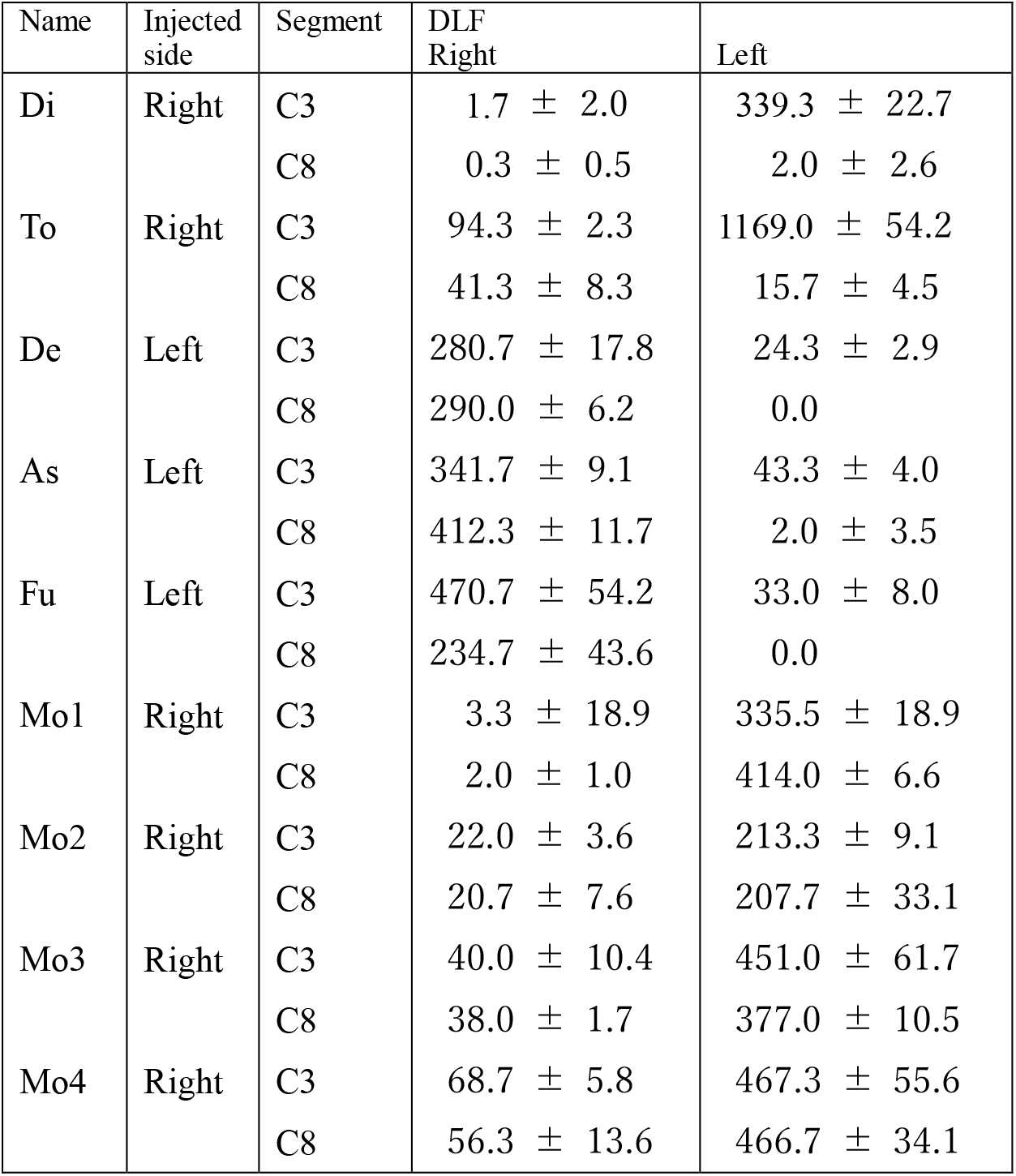
Mean ± SD of the number of labeled axons in the coronal slices in each component of the white matter at C3 and C8 in individual monkeys with coronal sections. The data from the intact animals were obtained from our previous study (Mo-1 to Mo-4 in Yoshino-Saito et al. 2010).

| Name | Injected side | Segment | DLF<br>Right | Left |
| --- | --- | --- | --- | --- |
| Di | Right | C3 | 1.7 ± 2.0 | 339.3 ± 22.7 |
|  |  | C8 | 0.3 ± 0.5 | 2.0 ± 2.6 |
| To | Right | C3 | 94.3 ± 2.3 | 1169.0 ± 54.2 |
|  |  | C8 | 41.3 ± 8.3 | 15.7 ± 4.5 |
| De | Left | C3 | 280.7 ± 17.8 | 24.3 ± 2.9 |
|  |  | C8 | 290.0 ± 6.2 | 0.0 |
| As | Left | C3 | 341.7 ± 9.1 | 43.3 ± 4.0 |
|  |  | C8 | 412.3 ± 11.7 | 2.0 ± 3.5 |
| Fu | Left | C3 | 470.7 ± 54.2 | 33.0 ± 8.0 |
|  |  | C8 | 234.7 ± 43.6 | 0.0 |
| Mo1 | Right | C3 | 3.3 ± 18.9 | 335.5 ± 18.9 |
|  |  | C8 | 2.0 ± 1.0 | 414.0 ± 6.6 |
| Mo2 | Right | C3 | 22.0 ± 3.6 | 213.3 ± 9.1 |
|  |  | C8 | 20.7 ± 7.6 | 207.7 ± 33.1 |
| Mo3 | Right | C3 | 40.0 ± 10.4 | 451.0 ± 61.7 |
|  |  | C8 | 38.0 ± 1.7 | 377.0 ± 10.5 |
| Mo4 | Right | C3 | 68.7 ± 5.8 | 467.3 ± 55.6 |
|  |  | C8 | 56.3 ± 13.6 | 466.7 ± 34.1 |

### The number of CST axon collaterals entering the gray matter

Figures 4A and B show transverse axons entering the gray matter (GM) in a coronal section of the C3 segment from the ipsilesional DLF in monkeys in which BDA was injected into the contralesional M1. The number of transverse axons among the total number of axons descending in the DLF at the C3 segment from the contralesional M1 was significantly larger in the lesioned monkeys than in the intact monkeys (Fig. 4C). Figures 4D and E show transverse axons entering the GM from the ipsilesional DLF in a longitudinal section. The number of transverse axons among the total number of axons descending in the ipsilesional DLF at the C3 segment was significantly larger in the lesioned monkeys than in the intact monkeys (Fig. 4F). Conversely, Figures 4G and H show transverse axons entering the gray matter of the C3 segment from the contralesional DLF in coronal sections of the monkeys in which BDA was injected into the ipsilesional M1. The number of transverse axons among the total number of axons descending in the contralesional DLF at the C3 segment was significantly larger in lesioned monkeys with BDA injection to the ipsilesional M1 than in intact monkeys (Fig. 4I). Transverse axons from the ipsilesional M1 tended to increase at the C8 segment, but the difference was not significant (Fig. 4J and K).

**Figure 4.**
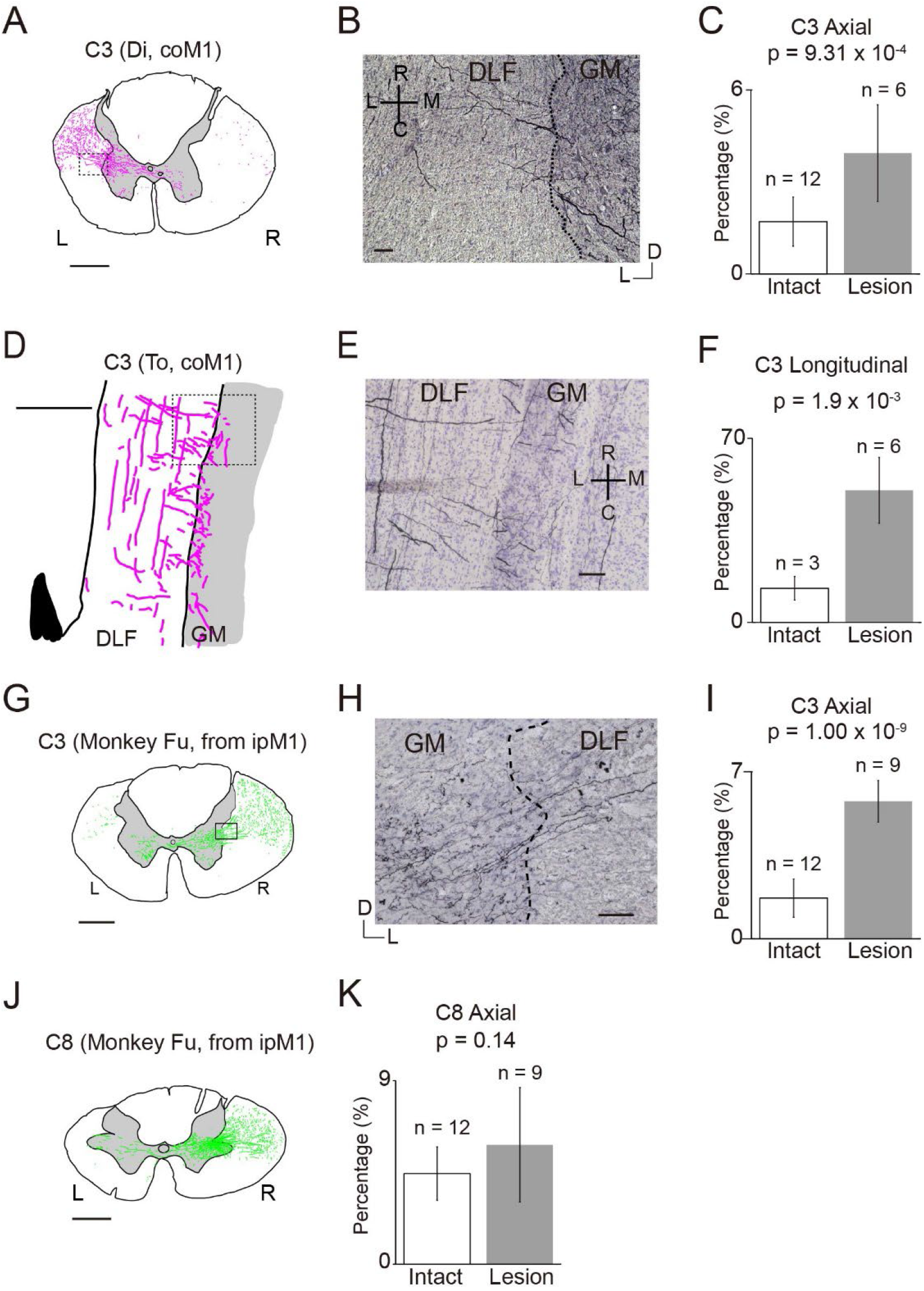
Collateral branching from CST axons originating from the contralesional M1 (coM1) or ipsilesional M1 (ipM1) at the C3 (**A-I**) and C8 segment (**J** and **K**). **A**. An example trace of CST axons in an axial section showing transverse CST axons entering the gray matter from the ipsilesional DLF at C3 in a monkey with BDA injection into the coM1 (Monkey To, the same as Fig.2D(ii), C3). Scale bar = 1 mm. **B**. A high magnification view of the square in **A**. The dashed line indicates the border between the gray matter and white matter (DLF). D, dorsal; L, lateral. Scale bar = 50 μm. **C**. Bar graphs indicate the percentage of transverse axons among the labeled axons in ipsilesional DLF at C3. Data were obtained by averaging the number from three axial sections of Monkeys As, De, and Fu. Error bars indicate standard deviation. I, intact monkeys (Mo-1,2,3,4); L, lesioned monkeys (Monkeys To and Di). **D**. An example trace of longitudinal section of the C3 segment showing transverse axons (black arrow) entering the gray matter from the ipsilesional DLF at C3 in a monkey with BDA injection into the coM1 (Monkey Gr). Descending axons are indicated with black arrowheads. A dashed line indicates the border between gray matter and white matter. R, rostral; C, caudal; L, lateral; Med, medial. Scale bar = 1 mm. **E**. High magnification view of the square in **D**. The dashed line indicates the border between gray matter and white matter. Scale bar = 100 μm. **F**. Percentage of transverse axons among the labeled axons in contralesional DLF at C3 in monkeys with BDA injection into the coM1 (Monkeys Al and Gr). **G**. An example trace of CST axons in an axial section showing transverse CST axons entering the gray matter from the contralesional DLF at C3 in the monkey with BDA injection into the ipM1 (Monkey Fu, the same as Fig.2D(iii), C3). Scale bar = 1 mm. **H**. High magnification of the square in **G**. Scale bar = 100 μm. **I**. The percentage of transverse axons out of the total number of labeled axons in the contralesional DLF at C3. Data were obtained from the axial sections of Monkeys Fu, As, and De. **J**. An example trace of CST axons in an axial section showing transverse CST axons entering the gray matter from the contralesional DLF at C8 in a monkey with BDA injection into the ipsilesional M1 (Monkey Fu, the same as Fig.2D(iii), C8). Scale bar = 1 mm. **K**. Percentage of transverse axons among the labeled axons in contralesional DLF at C8 in monkeys with BDA injection into the ipM1 (Monkeys De, As, and Fu).

### Axons descending in the gray matter bypassing the DLF lesion

In Figure 2C (ii) and Figure 3A, we described the existence of CST axons in the gray matter of the ipsilesional C8 segments originating from the contralesional M1. In the coronal sections obtained from Monkey To, we observed some axons that appeared to descend through the gray matter (GM) at the level of lesion (C4) and continued to descend in the gray matter to reach the C8 segments. We confirmed such descending axons in the longitudinal sections obtained from Monkeys Al (Fig. 5) and Gr. Figure 5A shows the overview of a sagittal section spanning from the C3 to C6 segment. Rostral to the lesions (the level between the two horizontal arrows in C4-C5), a number of descending axons in the white matter were sending collaterals to the gray matter, which terminated there (closed arrows in Fig. 5B). Many axons appeared to further run in the rostro-caudal direction in the GM. Figure 5C shows the stained CST axons in the GM caudal to the lesion (C5; between the two dotted lines in Fig. 5A) in Monkey Gr (Fig. 5C (i)) and in the corresponding area in the intact monkey (Fig. 5C (ii) from Mo-5 in Yoshino-Saito et al., (2010). Figure 5D (i) and (ii) analyzed the direction and length of individual axons in Figure 5C (i) and (ii), respectively, and the normalized results are shown in Figure 5E (i) and (ii). The results suggest that axons running in the longitudinal direction (close to 0º) already existed in the intact state. However, the proportion of long axons running in the longitudinal direction [Angle < 33.75º and Length > 500 μm in Fig. 5F (i), data from both Monkeys Al and Gr] appeared to have increased in the animals with SCI. Further, some of the longitudinal axons reached the C6/C7 segment and appeared to terminate on motoneuron-like large neurons (Fig. 5G and H). These results suggest that the number of CST axons descending in the gray matter increased after SCI and some of them established connections with motoneurons in the lower cervical segments.

**Figure 5.**
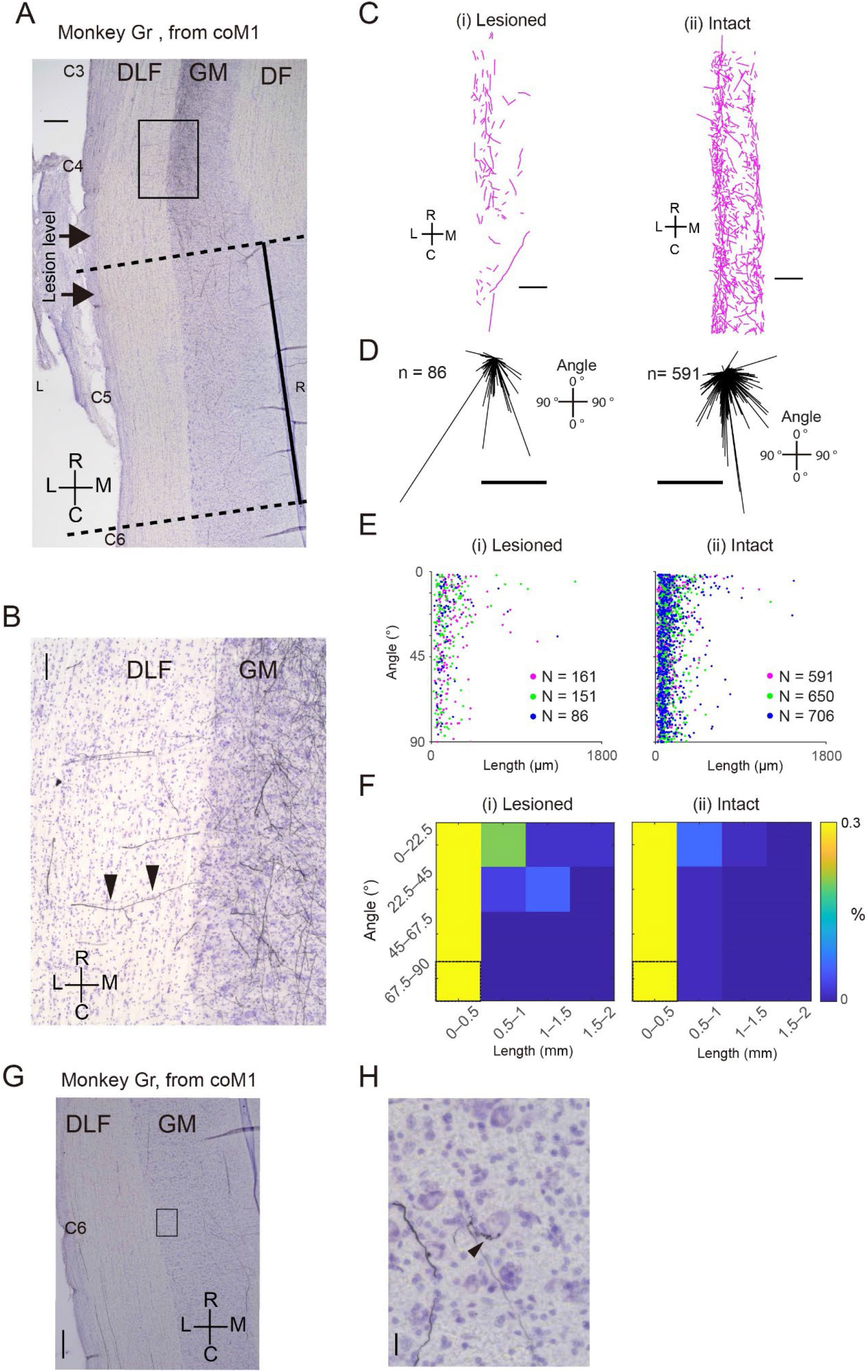
Longitudinal sections of the cervical spinal cord showing CST axons from the contralesional M1 (coM1) that descended in the ipsilesional gray matter. **A**. A photomicrograph of a longitudinal section spanning C3-C6 segments showing the BDA-labeled CST axons. Horizontal arrows at C4 – C5 indicate the levels of rostral and caudal borders of the C4/C5 lesion. The black square corresponds to the area of **B**. The dashed line indicates the rostral and caudal ends of **C**. The black line indicates the direction of the central canal, which corresponds to the R-C axis in **C**. Scale bar = 500 μm. **B**. A high magnification view of the square in **A**. Arrowheads show a BDA-labeled CST axon that ran horizontally and entered the gray matter from the DLF. Scale bar = 100 μm. **C**. Traced axons in the gray matter at the C5 segment of a lesioned monkey ((i); Monkey Gr) and intact monkey ((ii); Mo-5). Scale bars = 500 μm. **D**. Direction and length of individual axons in **C**(i) and **C**(ii) are indicated with the distribution of vectors in the left and right panels, respectively. Scale bar = 50 μm. **E**. Relationship between length of each axon piece and its direction shown in **D** in 3 sections from Monkey Gr (i) and Mo5 (ii) (different color assignment). **F**. Density map of the relationship between the length of each axon piece and its direction. Data obtained from **E**. The values were normalized by the overall number of the labeled axons in each condition. Encased bins indicate the baseline for the normalization in each panel. **G**. Photomicrograph of the C6 segment. Magnified view of the square is indicated in **H**, where the axons are terminated on a large, presumptive motoneuron. The black arrowhead indicates a presumptive button-like swelling terminating on the motoneuron. R, rostral; C, caudal; L, lateral; M, medial; DLF, dorsolateral funiculus; GM, gray matter; DF, dorsal funiculus. Scale bars in G and H indicate 500 μm and 20 μm, respectively.

### Distribution of CST axons and terminal buttons originating from the contralesional (right) M1

Figure 6 shows the relative distribution of axons and button-like swellings in each lamina (Fig. 2B) of the ipsilesional (left) and contralesional (right) C3 and C8 segments in the monkeys with BDA injections into the contralesional M1. The values were normalized as the percentage of descending axons in the DLF at the C3 segment on both sides. Rostral to the lesion (C3), as shown in Figure 6A, there was no significant difference in the number of axons in each lamina in both the contralesional and ipsilesional sides between the intact and lesioned animals. As for the number of button-like swellings, there was subtle increase in the ipsilesional lamina V and IX and contralesional lamina IX in the lesioned animals (Fig. 6B). Caudal to the lesion (C8), the number was drastically decreased due to the lesioning, however, some axons and buttons were confirmed in the lamina VI/VII and IX on the ipsilesional side (Fig. 6C and D), as has been described in the preceding sections (Fig. 2C (ii), 3A and 5C).

**Figure 6.**
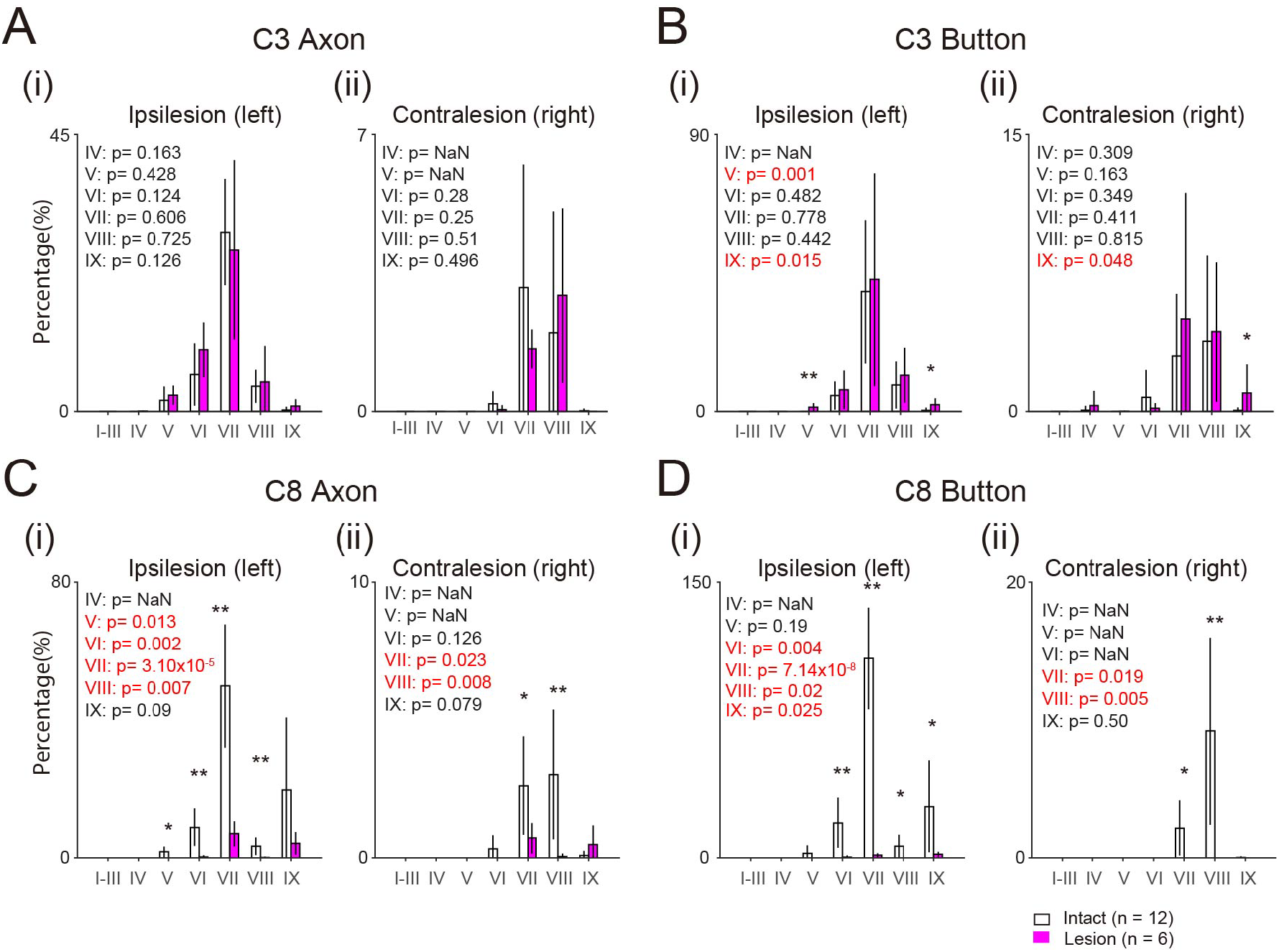
Comparison of the number of CST axons and buttons originating from the contralesional M1 between the intact and lesioned animals. **A**. Histograms indicate the number of axons in each lamina of the ipsilesional (i) and contralesional C3 (ii), divided by the total number of the axons in the DLF on both sides at C3. Insets in each panel indicate the p-value of t-test. **B**. The number of button-like swellings in each lamina of the ipsilesional (i) and contralesional C3 (ii). **C**. The number of axons in each lamina of the ipsilesional (i) and contralesional C8 (ii). **D**. The number of button-like swellings in each lamina of the ipsilesional (i) and contralesional C8 (ii). **, p<0.01; *, p<0.05. Red letters in the insets indicate statistically significant values.

### Distribution of CST axons and terminal buttons originating from the ipsilesional (left) M1

Figure 7 shows the relative distribution of axons and button-like swellings in each lamina (Fig. 2B) of the ipsilesional (left) and contralesional (right) C3 and C8 segments in the monkeys with BDA injections into the ipsilesional M1. The values were normalized as the percentage of descending axons in the DLF at the C3 segment on both sides. Rostral to the lesion (C3), as shown in Figure 7A, no significant difference was observed except for an increase in contralesional lamina VIII in the number of axons between the intact and lesioned animals. The number of button-like swellings decreased in the contralateral lamina VII in the lesioned animals (Fig. 7B). Caudal to the lesion (C8), the number of axons increased in laminae V, VI, VII, and IX on the ipsilesional side and in laminae V, VI, and VIII on the contralesional side (Fig. 7C). The increase in the axons on the ipsilesional side should be mostly derived from the crossing axons from the contralesional DLF, as shown in Fig. 3B. The number of button-like swellings also increased in laminae VI and IX, but decreased in lamina VIII on the ipsilesional side. Conversely, they also decreased in laminae VI/VII and IX on the contralesional side. Implication of these findings will be addressed in the Discussion.

**Figure 7.**
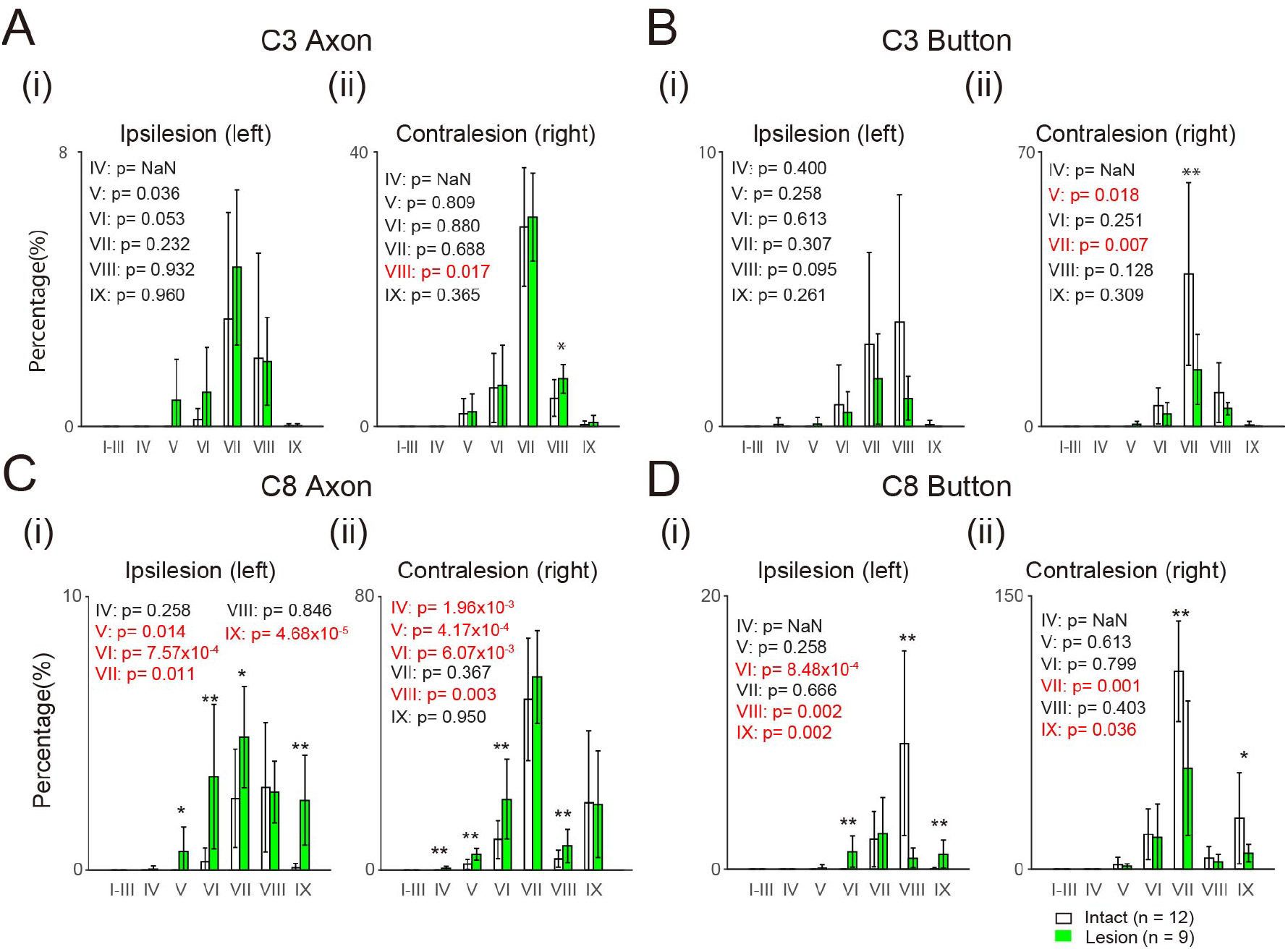
Comparison of the number of CST axons and buttons originating from the ipsilesional M1 between the intact and lesioned animals. **A**. Histograms indicate the number of axons in each lamina of the ipsilesional (i) and contralesional C3 (ii), divided by the total number of the axons in the DLF on both sides at C3. Insets in each panel indicate the p-value of t-test. **B**. The number of button-like swellings in each lamina of the ipsilesional (i) and contralesional C3 (ii). **C**. The number of axons in each lamina of the ipsilesional (i) and contralesional C8 (ii). **D**. The number of button-like swellings in each lamina of the ipsilesional (i) and contralesional C8 (ii). **, p < 0.01; *, p < 0.05. Red letters in the insets indicate statistically significant values.

## Discussion

In this study, we analyzed the trajectories of CST axons from both the contralesional and ipsilesional M1 in animals at 5-29 months after injury to clarify the neural pathways that support prominent recovery. Morphological changes in CST from the bilateral M1 were detected at spinal segments both rostral and caudal to the lesion. These changes included regeneration of the direct corticomotoneural connection from the contralesional M1 and emergence of those from the ipsilesional M1. Both phenomena already occurred in monkey Di sacrificed 5 months after the injury. The results suggest that the corticospinal pathways from bilateral M1 may mediate cortical commands to control hand motoneurons. Figure 8 summarizes the corticospinal projections before the lesion (Fig. 8A) (Yoshino-Saito et al. 2010) and after the full recovery (Fig. 8B). We further compared them with those reported previously on the hemisection models (Fig. 8C) (Galia and Darian-Smith, 1997; Rosenzweig et al. 2010).

**Figure 8.**
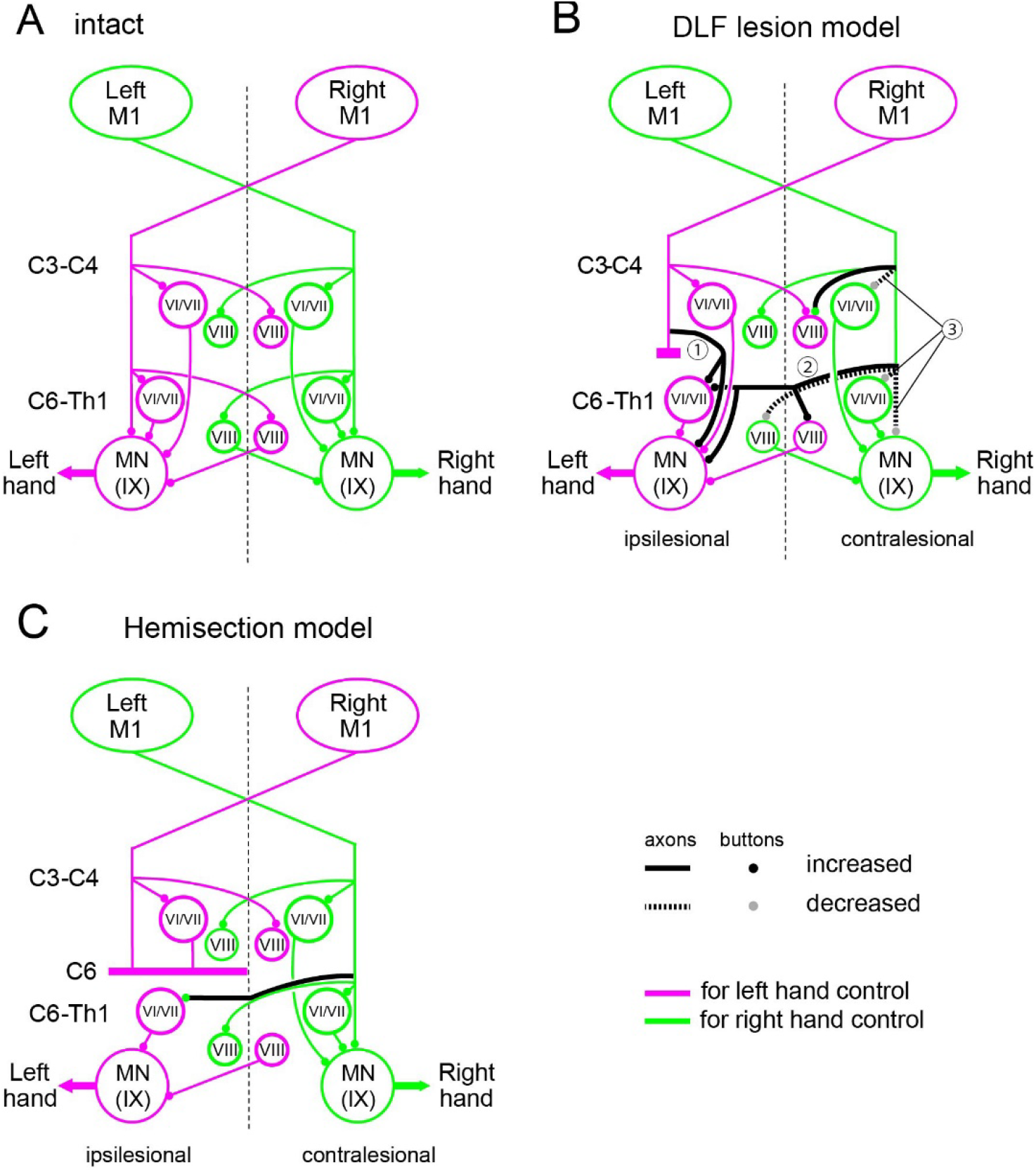
Schematic diagrams of the corticospinal projections originating from the forelimb area of M1 to individual spinal lamina of the C3-C4 and C6-Th1 segments in intact animals (**A**) and monkeys with the DLF lesion at C4/C5 (**B**). They are compared with the observation of the hemisection models by Galea and Darian-Smith (1997) and Rosenzweig et al. (2010) which showed poorer recovery (**C**). The magenta and green indicates the projections from M1 for left and right hand control, respectively. **A**. A schematic diagram of the projections of the CST projections from the bilateral M1 to the C3 and C8 segments summarized from our previous study (Yoshino-Saito et al. 2010). **B** and **C**. The same arrangement as **A**. Black lines and circles indicate the increased axons and buttons, respectively. Dotted black lines and gray circles indicate decreased axons and buttons, respectively. Uncrossed CST axons are intentionally removed from the figures because their contribution to the hand movement control has not been clearly demonstrated so far.

### Recovery mechanisms in the C4/C5 DLF lesion model

Previous studies have shown that the ability of dexterous hand movements can recover prominently in monkeys with DLF lesions that transected the l-CST and rubrospinal tract at the C4/C5 segment (Sasaki et al, 2004; Nishimura et al. 2007, 2009; Sugiyama et al. 2013; Sawada et al. 2015; Tohyama et al. 2017). In this model, blockade of C3-C4 PNs was shown to impair the recovering hand movements during the early recovery stage, but did not cause change during the late stage, suggesting involvement of other pathways in recovery (Tohyama et al. 2017). At the cortical level, reversible inactivation of the ipsilesional M1 impaired the recovering hand movements at the early stage, but did not in the late stage (Nishimura et al. 2007). These observations suggest that the long-term recovery is supported by different neural systems from the short-term recovery. As described above, in the short-term recovery, the residual descending pathways to the spinal interneurons including the C3-C4 PNs and their intersegmental connections to hand motoneurons would have been the major sources which compensated for the function of lesioned CST. In addition, the latent descending pathway from the ipsilesional M1 to motoneurons mediated by the reticulospinal neurons, which became evident after administration of 4-aminopyridine in cats (Jankowska et al. 2006), would be another source of early recovery. Thereafter, morphological changes in the residual CST might have happened and been involved in the long-term recovery. In present study, we focused on the morphological changes in corticospinal projections responsible for the long-term recovery (>3-4 months).

### CST projections from contralesional M1

Rostral to the lesion, collateral branches from the CST axons entering the gray matter of the C3 segment from both the contralesional and ipsilesional M1 were increased (Fig. 4A-I). However, no marked change was observed in the number of axons and button-like swellings in the gray matter (Figs. 6A, B and 7A, B). The increase of the collateral branches may suggest that the PNs in the C3 segment might contribute to recovery at some stage after the injury. However, lack of increase in the axons and buttons in the C3 gray matter suggests that they were no longer playing a major role or functioning at best to the similar extent as the intact animals for the long-term recovery. This is consistent with our previous results, in which selective blockade of the PNs caused clear deficits during the early recovery but it was not the case long after the lesion (Tohyama et al. 2017). Caudal to the lesion (C8), axons (Fig. 6C) and button-like swellings (Fig. 6D) originating from the contralesional M1 decreased drastically by the lesion. However, some axons and terminals were observed in laminae VI/VII and IX (Fig. 6C and D) which might have originated, at least in part, from recrossing of uncrossed CST axons descending in the contralesional DLF. However, our microscopic investigation suggests that the number of recrossing axons (Fig. 3A) was not sufficient to explain the majority of axons and buttons. Instead, we confirmed that the majority of these axons were derived from the CST axons which originated from the ipsilesional DLF, projected to the gray matter rostral to the lesion, and descended in the gray matter bypassing the lesion (Figs. 3A, 5F)(➀ in Fig.8B). Such axons were observed in both Monkeys Di and To at 8 months (233 days) and 22.5 months (677 days) after injury, respectively. Comparison with intact animals suggested that some of these descending axons that bypassed the lesion would have emerged after the SCI (Fig. 5). Emergence of similar axons was reported in animals after the subhemisection at C6 by the administration of antibody against MAG which is released from glial cells and prevents neural regeneration (Nakawaga et al. 2015). The present study revealed that the long distance generation of axon branches spanning the C4/C5 segment to the C8 segment could have occurred spontaneously in the long-term by rehabilitative training.

### CST projections from ipsilesional M1

On the other hand, there was also a considerable increase in axons and button-like swellings originating from the ipsilesional M1, in the ipsilesional gray matter of the C8 segments spanning laminae V, VI, VII, and IX (Fig. 7C)(➁ in Fig.8B). Axons in the contralesional gray matter also increased, but the buttons decreased in the ipsilesional lamina VIII and contralesional laminae VII and IX (➂ in Fig.8B). The normal CST projections are directed to the contralateral VI/VII and IX and ipsilateral lamina VIII after recrossing (Kuypers 1982; Armand et al., 1997). Previous studies revealed spinal commissural neurons in the lamina VIII, which cross the midline to the contralateral side, terminate on interneurons in lamina VII or motoneurons in lamina IX, and play critical roles in left-right coordination (Silos-Santiago and Snider, 1992; Eide et al., 1999; Stokke et al., 2002; Matsuyama et al. 2006; Abbinanti et al., 2012). Therefore, the decrease in the number of terminals in laminae VI/VII and IX on the contralesional side and in lamina VIII on the ipsilesional side (➂ in Fig.8B), and the increase in the number of axons in laminae V, VI, VII, and IX and that in number of terminals in laminae VI and IX on the ipsilesional C8 (➁ in Fig.8B) suggest that the descending control by the ipsilesional M1 might have partly switched from control of the contralesional hand to that of the ipsilesional hand. Behaviorally, we did not observe any decline in the dexterity of the contralesional (intact side) hand movements, presumably because of redundancy in the intact motor system. The mechanism of such switch is unclear, but should be an interesting target of future studies. All these observations were observed in the monkey Di at 5 months after the SCI and summarized in Fig. 8B.

### Comparison with hemisection/subhemisection model

It is important to compare the results in the monkeys with C4/C5 DLF lesion that showed prominent recovery to the monkeys with larger lesions with poorer recovery (Galea and Darian-Smith, 1997; Rosenzweig et al. 2010). In the hemisection model, extension of the recrossing CST axons from those descending in the contralesional side was found, but changes in the other components were not described. On the other hand, in the subhemisection model by Nakagawa et al. (2015) treated with anti-MAG antibody, the CST axons were extended, descended in the spinal gray matter bypassing the lesion and were connected to the motor nuclei (like Figure 8B➀). These animals showed prominent recovery in grasping. Thus, reconnection of the CST axons to the motoneurons might be one of the critical components for prominent recovery of hand dexterity.

After the full recovery of precision grip, the contralesional M1 retained the axons and terminals rostral to the lesion, presumably involving the control of C3-C4 PNs, and increased the direct control of caudal segments by extending axons that descend in the gray matter. In addition, the ipsilesional M1 increased recrossing axons caudal to the lesion, suggesting shift of its control from the intact hand to the affected hand. Thus, although previous studies suggest that the contribution of C3-C4 PNs (Tohyama et al. 2017) and the ipsilesional M1 (Nishimura et al. 2007) may decrease in the late stage of recovery, this does not always indicate that their role was completely replaced. Instead, CSTs originating from both the contra- and ipsilesional M1 still contribute to recovery after the SCI through mono- and/or multi-synaptic pathways to motoneurons (Ninomiya et al. 2022), as suggested by massive reorganization of cortical networks (Nishimura et al. 2007; Nardone et al. 2013; Suzuki et al. 2020. Thus, the brain distributes its control over a variety of neural systems for long-term recovery. This strategy may assure robustness of motor control in the brain with a partial lesion, to avoid fatal damage induced by additional injury. Furthermore, in this study, we focused on the change in the axonal trajectories and terminations of the CST. However, it is highly likely that other descending motor systems such as rubrospinal and reticulospinal tracts contribute to long-term recovery, which should be the subject of future studies.

## Acknowledgements

We thank Y. Itani, N. Takahashi, M. Togawa, and K. Isa for technical help. The monkeys used in this study were provided by the National Institute of Natural Sciences through the National Bio-resource Project of the Ministry of Education, Culture, Sports, Science and Technology of Japan (MEXT). This work was supported by grants from the “Brain Machine Interface Development” and performed under the Strategic Research Program for Brain Sciences from MEXT and the Japan Agency for Medical Research and Development to Y.N., a Grant-in-Aid for Scientific Research on Innovative Areas “Adaptive circuit shift” to T.I. (Project no. 26112008) and H.O. (26112003), and a Grant-in-Aid for Scientific Research from MEXT to H.O. [KAKENHI (B) no. 24300196 and (A) no.15H01819]. All data are stored at the Department of Developmental Physiology at the National Institutes for Physiological Sciences, Okazaki, Japan.

